# Yeast cells under glucose-limitation environment need increased response cost for osmostress defense

**DOI:** 10.1101/2021.06.24.449709

**Authors:** Wenting Shen, Ziqing Gao, Kaiyue Chen, Qi Ouyang, Chunxiong Luo

## Abstract

Cells always make responses to environmental changes, involving dynamic expression of tens to hundreds of proteins. This response system may demand substantial cost and thus affect cell growth. Here, we studied the cell’s responses to osmostress under glucose-limitation environments. Through analyzed thirteen osmotic-downstream proteins and two related transcription factors, we found that the cells required stronger responses under low glucose concentrations than normal glucose condition after being stimulated by osmostress, even the cell growth rate was unchanged in these two constant conditions. We proposed and verified that under a glucose-limitation environment, the glycolysis intermediates were limited (defense reserve saving), which caused that cells needed more glycerol production enzymes to adapt to the osmostress. Further experiments proved that this ‘defense reserve-saving’ strategy required cells to spend more response cost when facing stress, which on the other hand, enhanced the fitness for the coming environment variations via protein accumulation reserve.

## Introduction

Organisms need to sense, respond, and adapt to environments for survival and growth with optimized strategies for growth competitions. As examples, bacteria carry out chemotaxis to avoid unfavorable conditions and migrate toward favorable locations(Adler, 1966)(Zhang *et al*, 2019); *Saccharomyces cerevisiae* (budding yeast) cells orchestrate a common gene expression program called the environmental stress response (ESR) to adapt to environmental changes(Gutin *et al*, 2015a). Besides, cells also change the proteome or energy allocation under different growth rates or growth conditions(Boer *et al*, 2003). For bacterial cells under steady-state exponential growth, it is well known that the proteome organization predominantly depends on the growth rate restricted by the nutrition quality of the medium(Scott *et al*, 2010). As the growth rate decreases in nutrition limitation environment, the cell tends to invest more proteomic resources into intake systems rather than translation and metabolism(Mori *et al*, 2016). However, allocation of limited resources and stress response are always considered independently in previous work. The strategy of cells preparing and responding to stress under different nutrients environments is still to be determined.

Moreover, even if cellular physiology such as growth rate remains constant, the proteome partitioning and energy allocation may also have discrepancies. Previous studies have shown that in *Saccharomyces cerevisiae*, despite the availability of poor nutrient sources, cell growth rate under low glucose concentration (0.1%) is similar to normal condition (2%)(Youk & Van Oudenaarden, 2009). However, translation and metabolism have changed even under these constant environmental conditions. Genes involved in uptake glucose and carbohydrate-related metabolism have a higher expression level at 0.1%(Chen *et al*, 2020a). To achieve optimal growth, cells might choose efficient pathways and abandon inefficient investments. Since growth and stress defense always compete for limited resources, it will be intriguing to know what stress defense strategy will the cell adopt under the low glucose condition(0.1%) compared with rich glucose condition(2%)(Ho & Gasch, 2015)?

In the natural environment of yeast, the most common stress is the rapidly changing water activity(Hohmann, 2002). Thus, cells are often exposed to hyperosmostress and need to maintain water balance. The osmoregulatory system in the yeast is quite well understood. The osmo-adaptation network integrates gene expression and regulation of metabolic flux. The key to yeast osmoregulation is the production and accumulation of the compatible solute glycerol, which is mainly regulated by HOG mitogen-activated protein kinase (MAPK) pathway(Dihazi *et al*, 2004)(Hohmann, 2015). Specifically, the redistribution of glycolytic flux from growth to glycerol production is the main process of osmoadaptation(Bouwman *et al*, 2011)(Petelenz-Kurdziel *et al*, 2013). Therefore, in this work, we mainly discussed the stress response and the defense strategy of budding yeast to the osmostress under different glucose concentrations (Fig. 1A).

**Figure 1.**
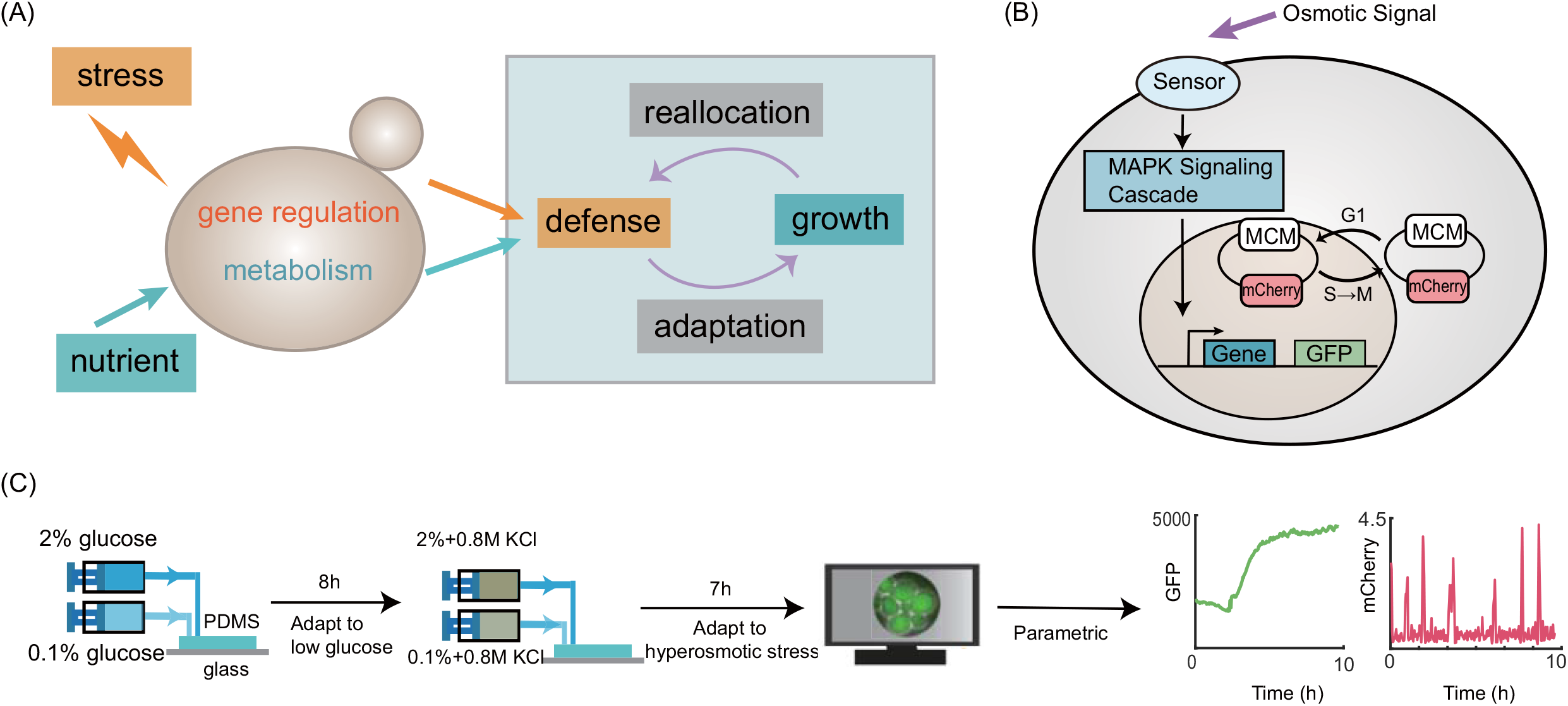
Schematic model, strain background and experimental settings. A. A simplified view of the trade-off between growth and stress defense. Nutritional restriction affects growth rate through metabolism and gene regulation, while stress response and growth compete for limited resources. We wonder if in a nutrient limitation environment where the growth rate does change significantly, would the stress response behave differently. B. Schematics for monitoring target protein dynamics (with fluorescent protein fusions) and cell cycle (with the MCM system) in single cells. C. Yeast cells are cultured independently under 2% and 0.1% (m/V) glucose for 5 hours, and then exposed to hyperosmotic stress (0.8M KCl). The adaptation process was observed under microscope.

For more details, we studied single yeast cell’s protein dynamics to hyperosmostress under glucose limitation environment using a microfluidic device. The synthesis rates of downstream proteins and the growth rate of cells were measured. It was found that cells had higher synthesis rate of osmotic-related proteins when facing same osmostress at lower glucose conditions. Moreover, a similar phenomenon was found in the nuclear localization behavior of transcription factor Hog1. Given the important role of glucose in metabolic during osmoadaptation, we speculated that under a low glucose environment with limited glycolytic flux, cells required more glycerol production enzymes to adapt to hyperosmostress. A simplified model based on this assumption and the experimental data was established to explain the osmotic protein dynamics. The model also helps to clarify of resource allocation strategies in glucose-limited environments. That is, yeast reduces the metabolic reserve for optimal growth under the condition of continuous limited glucose, which is called the “defense reserve saving”. Therefore, yeast needs to invest more to synthesize related proteins when facing stress, which corresponds to higher response costs. Further experiments showed that the accumulation of a large amount of stress-related proteins under low glucose condition can improve the fitness of cells to the second osmotic-stimuli. This resource reallocation of “defense reserve saving” may facilitate our understanding of the operation and the design of complex biological systems.

## Results

### The stress response of cells to osmostress under different glucose concentration conditions shows obvious differences

To characterize the stress response of budding yeast in a glucose limitation environment, we monitored yeast cells with GFP-tagged stress-responsive proteins using a microfluidic system (Fig. S1). The strain also has protein MCM labeled by m-Cherry as a cell cycle marker (Fig. 1B). MCM complex is imported into the nucleus in G1 phase and exported from the nucleus during the S phase, thus it can be used to easily quantify the durations of cell cycles (Chen *et al*, 2020b)(Braun & Breeden, 2007)(Fig. S2).

For a low glucose environment, we chose a glucose concentration of 0.1% to ensure that the growth rate is not significantly affected(Chatterjee & Acar, 2018). Cells were initially cultured for 8 hours in a microfluidic system with a glucose concentration of 2% or 0.1% to adapt to the environment, at that time the expression level of stress-responsive proteins and the duration of cell cycle remain constant. The duration of cell cycle was measured to be similar about 1.25 hours under 2% and 0.1% glucose conditions (for mother cells, Fig. S3). Then, cells were exposed to osmostress (0.8M KCl, the same for glucose concentration of 2% and 0.1%) for 7 hours (Fig. 1C). Bright field and fluorescence images were collected every 2.5 min for dynamic studies.

When exposed to osmostress, the osmotic stress proteins such as GPP1, a well-characterized Hog-MAPK downstream protein(Påhlman *et al*, 2001), were upregulated in both 0.1% and 2% glucose conditions (Fig. 2A). Strikingly, it is observed that higher expression level of Gpp1-GFP under 0.1% glucose condition than 2% glucose. Meanwhile, the cell cycle went through a phase of shock (Fig. S4), because the extended G1 phase could help to resist adverse environments(Yin *et al*, 2003). While under glucose limitation conditions, cells seem to employ a longer recovery time. After the first generation to resume growth, the cell cycle gradually almost adapted to the initial level (Fig. 2B). On account of the similar steady growth rate at 0.1% and 2% glucose, we supposed that cells could efficiently adapt to hyperosmotic under glucose limitation condition (0.1%).

**Figure 2.**
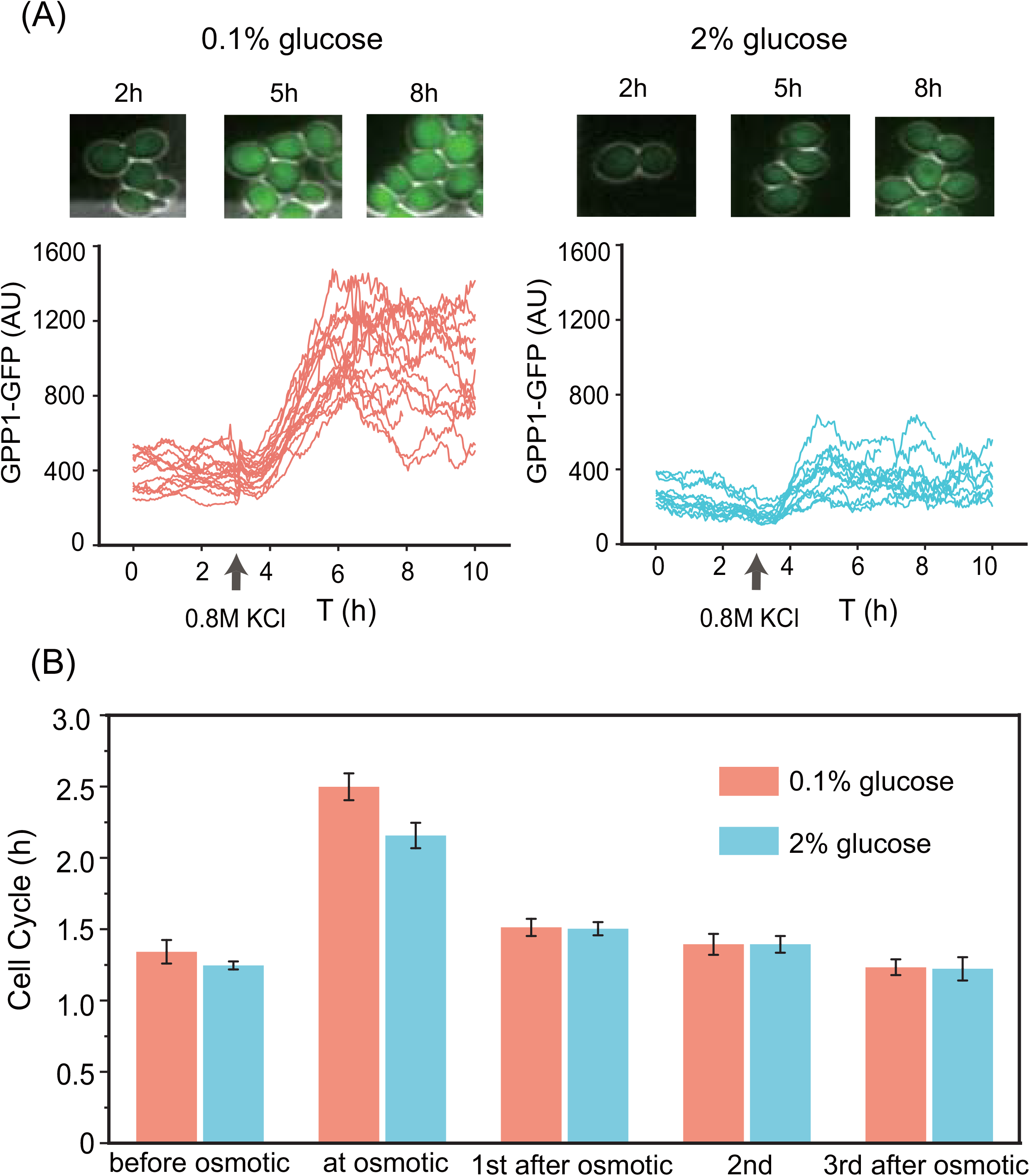
Preliminary phenomena of osmotic response under different glucose concentrations. A. Time-lapse microscopy of Gpp1-GFP in response to 0.8 M KCl. Examples of raw images (top) and traces of individual cells (bottom). The left is the low glucose (0.1%) condition, and the right is under normal glucose environment (2%). B. Histogram of single cell growth rate at different glucose concentrations. Numbers of cells in 2%, and 0.1% glucose concentrations are 80 and 89, respectively. Error bars represent SEM.

The osmotic adaptation in a rich medium (2% glucose) has been extensively studied (Hohmann, 2002). Interestingly, in a low-glucose environment, stress-related proteins show a more pronounced response to osmostress, while the growth rate does not seem to be restricted. We were curious about the mechanisms of the dramatic stress response differences between 0.1% glucose condition and 2%. This implies that although cellular physiology was not affected, resource reallocation may have occurred in a glucose-limited environment.

### Stress response proteins have a higher synthesis rate in a glucose-limited environment

To systematically study this phenomenon, we measured the dynamics of proteins in response to osmostress, including 5 osmotic induced proteins, 4 general stress-responsive proteins, and 4 proteins related to other stress. (Table S1). We observed that the level of target proteins at different glucose concentrations showed no significant difference under constant nutrient supply. Then after switching to a hyperosmotic environment, the stress-responsive proteins were upregulated and exhibited an obvious difference in different glucose environments (Fig. 3A, Fig. S5).

**Figure 3.**
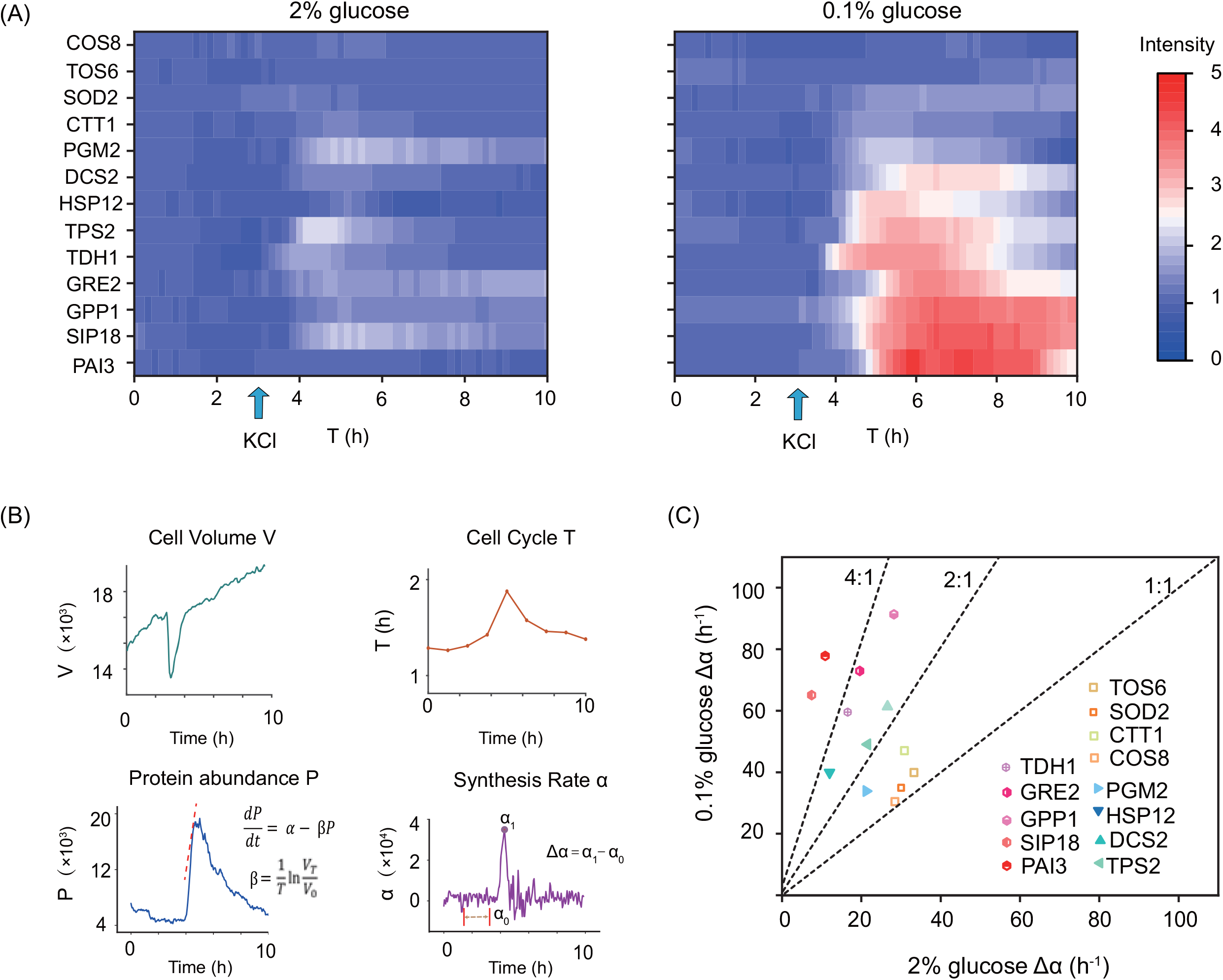
Modulation of the protein dynamics to osmostress. A. Protein dynamics at different glucose concentrations. 0.8M KCl was added at 3h. Data was normalized with the initial 2 hours of 2% glucose, with greater expression indicated by red and less by blue. B. Parametric decomposition of response curves, from protein abundance and dilution rate to get the synthesis rate. Stress response cost Δαis the maximum synthesis rate after hyperosmotic minus the average synthesis rate before stimulation (1~2 hours). C. Δαat different conditions. Δαof osmotic response proteins at 0.1% glucose exceeds twice that of 2% glucose (red hexagon). General stress response proteins are marked as blue triangles. Points close to the 1:1 line are other functional proteins (yellow square).

Protein synthesis rate was used to characterize these responses, which was described by the following equation:

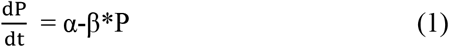

where P demoted the protein concentration, α and β respectively denoted the protein synthesis rate and the decay rate. The decay rate β is mainly determined by dilution, as the protein degradation under fast response can be neglected(Gutin *et al*, 2019). Thus, the maximum synthesis rate difference Δα between after stress and before stress could be calculated from time-course data, which reflects the stress response cost (Fig. 3B).

We noticed that most of the stress-related proteins had higher expression levels when facing osmostress under a low glucose environment (Fig. 3C). Among them, the Δα of Hog-MAPK pathway related proteins under low glucose is more than twice that of the normal condition (protein PAI3, SIP18, GPP1, GRE2, TDH1 marked as a red hexagon in Fig. 3C), while Δα of other proteins (protein TOS6, SOD2, CTT1, COS8, PGM2, HSP12, DCS2, TPS2, which are not directly influenced by Hog-MAPK pathway) show fewer differences when facing osmostress wherever in low glucose or normal conditions.

### Response dynamics are indicated by the transcription level

Having observed the significant increase in stress response to osmostress at 0.1% glucose compared with the normal 2% glucose condition, we wondered whether these different dynamical profiles were associated with transcription factors. In our description of the response process, protein synthesis correlated with the integral of mRNA, which was proportional to the integral of transcription factor activity (Hao & O’Shea, 2012).

We measured the relative nuclear intensity of Hog1, the essential transcription factor for initiating gene expression during osmoregulation of the budding yeast. Although single cells exhibited variability in nuclear localization, Hog1 shows a higher nuclear concentration in a glucose limitation environment when exposed to osmostress. Besides osmotic-specific transcription regulation, we also monitored the general response transcription factor Msn2, which is mainly controlled by PKA pathway. Different from Hog1, the nuclear localization traces of Msn2 exhibit no obvious difference between different glucose concentrations (Fig. 4A).

**Figure 4.**
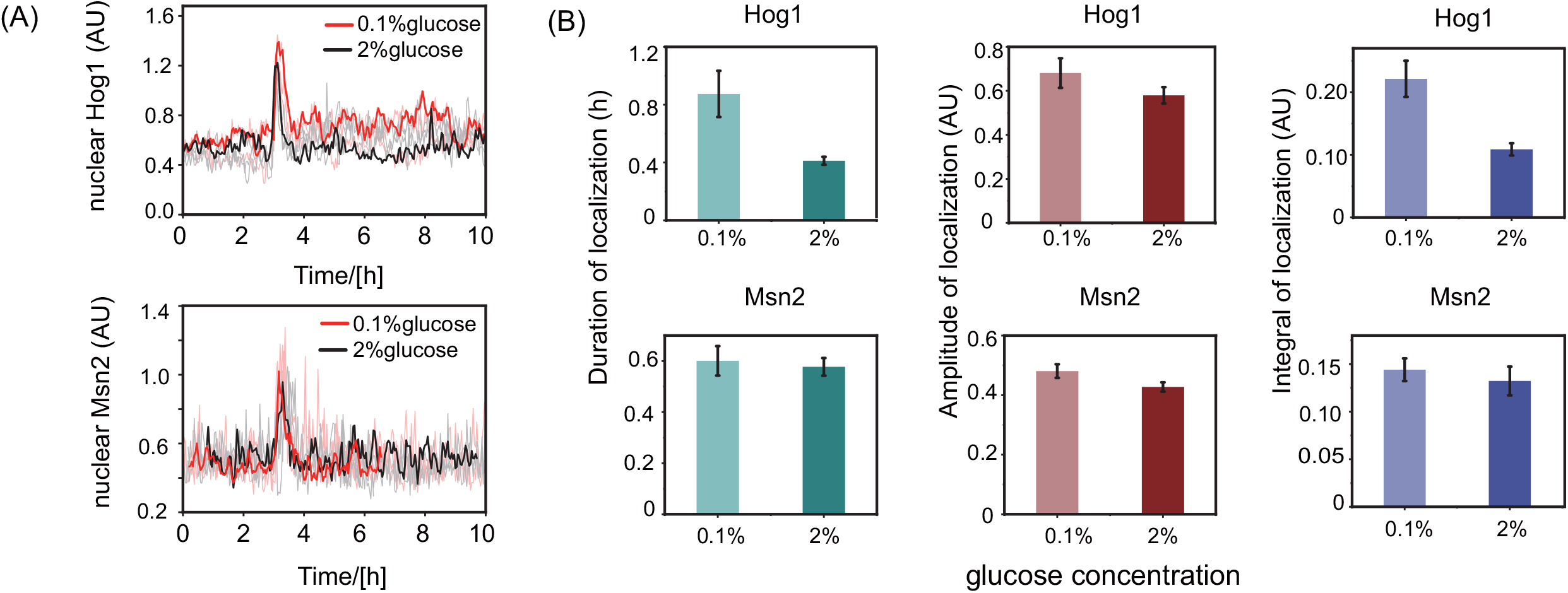
Transcriptional regulation in population of cells. A. Localization trajectory of single cells. The bold line is the single cell whose nuclear integral is the median value in the population. B. Duration, amplitude and integral of Hog1 and Msn2 nuclear localization are quantified. Error bars represent SEM. Table S2 lists the P values resulting from comparing duration, amplitude and integral of Msn2 and Hog1 nuclear localization between 2% and 0.1% glucose concentrations.

Then we quantified the dynamic characteristics of Hog1 and Msn2 localization in every single cell and calculated the average values across the populations in different glucose concentrations (duration, amplitude, and integral area; Fig. 4B). Expectedly, we found that the duration and integral of Hog1 nuclear localization significantly increase under glucose-limitation conditions (Fig. 4, Fig. S6). While the integrals of Msn2 nuclear localization show little difference between 2% and 0.1% glucose conditions.

Taken together, these results suggest that the nuclear localization of both Msn2 and Hog1 are required to resist osmostress. But cells tend to allocate more resources to the Hog pathway in a glucose-limited environment.

### Metabolic reconfiguration strategy for *S. cerevisiae*

When facing osmostress, yeast cells produced more osmotic-related protein and required a longer recovery time under 0.1% glucose condition compared with normal 2% glucose condition. To explain this higher response cost, we considered the metabolic reconfiguration for osmotic adaptation at different glucose concentrations.

Glucose is used by several cross-linking pathways once entering the cell. The main route of glucose utilization is glycolysis, which converts glucose to trehalose, glycogen and glycerol, etc (Teusink *et al*, 2000). The most important branch in unstressed cells is the carbon flux to biomass for growth. When facing osmostress, the cells will re-distribute the glycolytic flux from growth to glycerol production for osmotic balance. Limited glucose may affect the metabolic flux rerouting to glycerol accumulation through its intermediate metabolism product glycerol-3-phosphate (G3P) (Fig. 5A).

**Figure 5.**
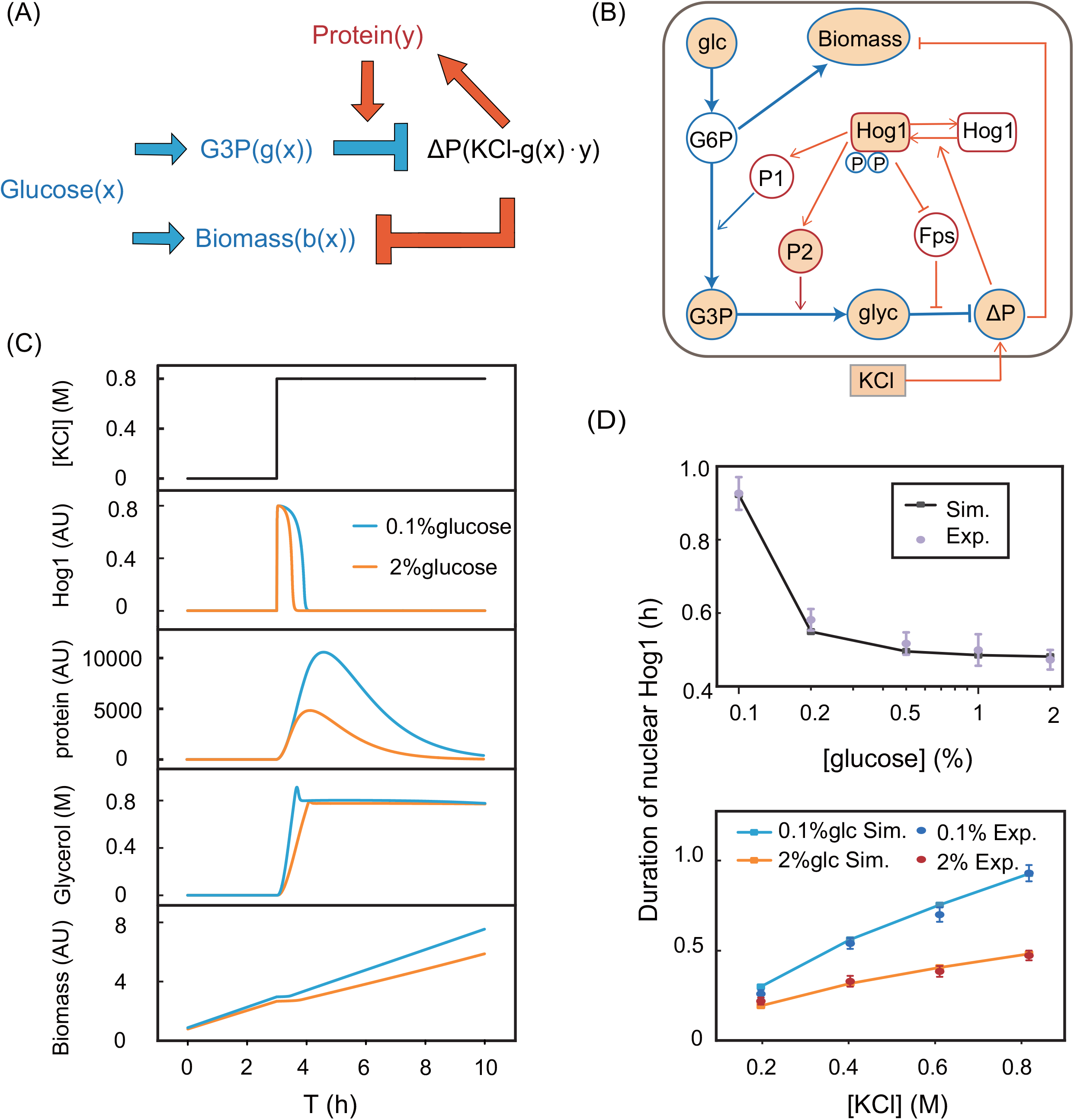
The simplified model can explain enhanced stress response under glucose-limitation condition. A. A schematic of the interplay between the osmotic adaptation and metabolic reconfiguration. Osmoadaptation requires compatible solute glycerol to compensate for a given stimulus. The core network structures describes the rerouting of glycolytic flux from growth to glycerol. The theoretical explanation for the experimental phenomenon is that lower glycolytic flux can maintain the normal growth of cells, but the reduction of the intermediate product G3P requires more glycerol synthase to synthesize a sufficient concentration of glycerol. B. Diagram of major signaling pathways involved in osmoadaption. Blue indicates glycolysis modulation. Orange indicates gene regulation. Measured entities are highlighted yellow. C. Model simulation for phosphorylated Hog1, downstream protein, intracellular glycerol and biomass synthesis. D. Duration of nuclear Hog1 reflects the adaptation time. The dots with error bars are experiment results. Error bars from three independent experiments. The dots with lines are model predictions.

Our experimental results showed that in response to hyperosmotic, the glycolysis rerouted to G3P in 0.1% glucose concentration was much less than that in 2% glucose (Fig. S7). Previous studies also indicated that the intracellular concentration of G3P decreased with external glucose concentration(Cronwright *et al*, 2002). Since the substrate for glycerol synthesis was reduced in a low glucose environment, more enzymes were required to accumulate glycerol to balance the osmotic pressure (Fig. S8A).

Thus, the emerging picture is that, under hyper-osmotic stress, the process of osmoadaptation leads to a coordinated activation of the environmental stress response and suppression of biomass formation. Δα indicates the protein cost, cell cycle arrest represents the glycolysis reallocation expense. Therefore, when nutrients are poor for redirecting sufficient glycolytic flux, cells will allocate more resources to stress-responsive gene expression, which means higher response costs.

### A simple model for metabolic-combined osmotic adaptation

To corroborate the above hypothesis in *silico*, we developed a computational model including the main metabolic features and feedback mechanisms of the osmotic adaptation. Specifically, glucose-6-phosphate (G6P) is the main product of glucose phosphorylation, which in turn converts to biomass and G3P. The glycolytic flux to G3P could be regulated via Hog1-mediated activation of phosphofructose-2-kinase, denoted by protein P1. Then, G3P is transformed to glycerol by the G3P phosphatases, represented by protein P2 (Påhlman *et al*, 2001)(Norbeck *et al*, 1996). Excessive glycerol leaks out through the glycerol facilitator Fps1, which appears to be controlled by Hog1 (Lee *et al*, 2013). On the other hand, we assumed that Hog1 phosphorylation and the reduction in biomass production were influenced by osmotic pressure discrepancy. The schematic of the model is illustrated in Fig.5B, and the details of coupled ordinary differential equations are included in supplemental data.

The simulated time traces of the model are shown in Fig. 5C. In response to osmostress, cells in a sufficient nutrient environment reallocated enough glycolytic flux, and had a faster glycerol production rate. While the glucose limitation environment required a longer nuclear duration of Hog1 to finally adapt. To examine whether this model can describe Hog1 dynamics, we also compared the experimental results with the simulation results in more conditions (Fig. 5D). Consistent with the model prediction, the duration of Hog1 nuclear localization increased with stress intensity upon glucose limitation and osmostress. The previous models focused primarily on describing the Hog-MAPK signaling pathway, but our model is capable of reproducing the Hog1 dynamics and cell physiology under varying levels of glucose concentration.

Therefore, this simplified model suggests that the metabolic intermediate for glycerol production in a glucose limitation environment is insufficient, thus more stress-related proteins are required. A previous study reported that at normal glucose condition, yeast stored excess carbons in several metabolic pathways, and the first step of osmostress response was to route internal stores in metabolic flux to glycerol production, accelerating osmo-adaptation(Bonny *et al*, 2021). While in glucose-limited conditions, if cells still maintain the metabolic reserve for defense, their instantaneous biosynthesis would be reduced? and adversely affect growth. Indeed, slow-growing cells are better prepared for stress than fast-growing cells(Zakrzewska *et al*, 2011), which may make it a good strategy. However, our results showed that the growth rate of cells under 0.1% glucose was almost the same as at 2%. In other words, cells maximized the growth rate, which means that the defense reserve will inevitably be reduced. Moreover, previous research suggests that yeast will reduce the reserves of the metabolite G6P when the glucose is reduced(Cronwright *et al*, 2002). This indicates that despite the ostensible advantage of faster adaptation, there is a trade-off between the biomass increase and the preparatory to subsequent insults.

Our results assumed that yeasts selected different metabolism pre-allocation strategies at different nutritional conditions. For cells under a glucose limitation environment, defense reserve savings provide current growth benefits at a cost of the recovery processes of osmo-adaptation.

### Enhanced stress response for defense

However, in osmo-adapted cells, the synthesis of downstream protein is overexpressed for glycerol production, which is more pronounced when glucose is limited. To explore the potential benefits of enhanced osmotic response under a glucose limitation environment, we monitored the cell growth rate under dynamically controlled osmolarity. Consecutive stress time (1 h) guaranteed sufficient glycerol synthesis. The first and second osmotic stress were separated by 1.5h stress-free medium (about one cell cycle). Then, the response cost of two stimulation can be compared. Our model predicted that cells could adapt to the subsequent osmostress more quickly due to the accumulation of downstream proteins during priming stimulus. Correspondingly, the translocation duration of Hog1 will decrease. This is consistent with the cellular memory theory(Ben Meriem *et al*, 2019).

Still using GPP1 as the reporter, we observed that after the first stimuli, Gpp1 was not diluted to the initial value both in 2% and 0.1% glucose condition (Fig. 6B). In response to the second stimulation under the condition of 0.1% glucose, the initial level of GPP1 was much higher than 2% glucose. In addition, the cell cycle arrest time at the second stimulation was reduced compared with the first stimulation (Fig. 6C). Especially, in 0.1% glucose condition, the arrest time at the second stimulation is basically the same as 2% glucose condition, while at the first stimulation it is longer than that under the condition of 2% glucose. This may because much more protein was accumulated for the rapid glycerol synthesis, and cells in a low-glucose environment can also resume growth as the normal condition.

**Figure 6.**
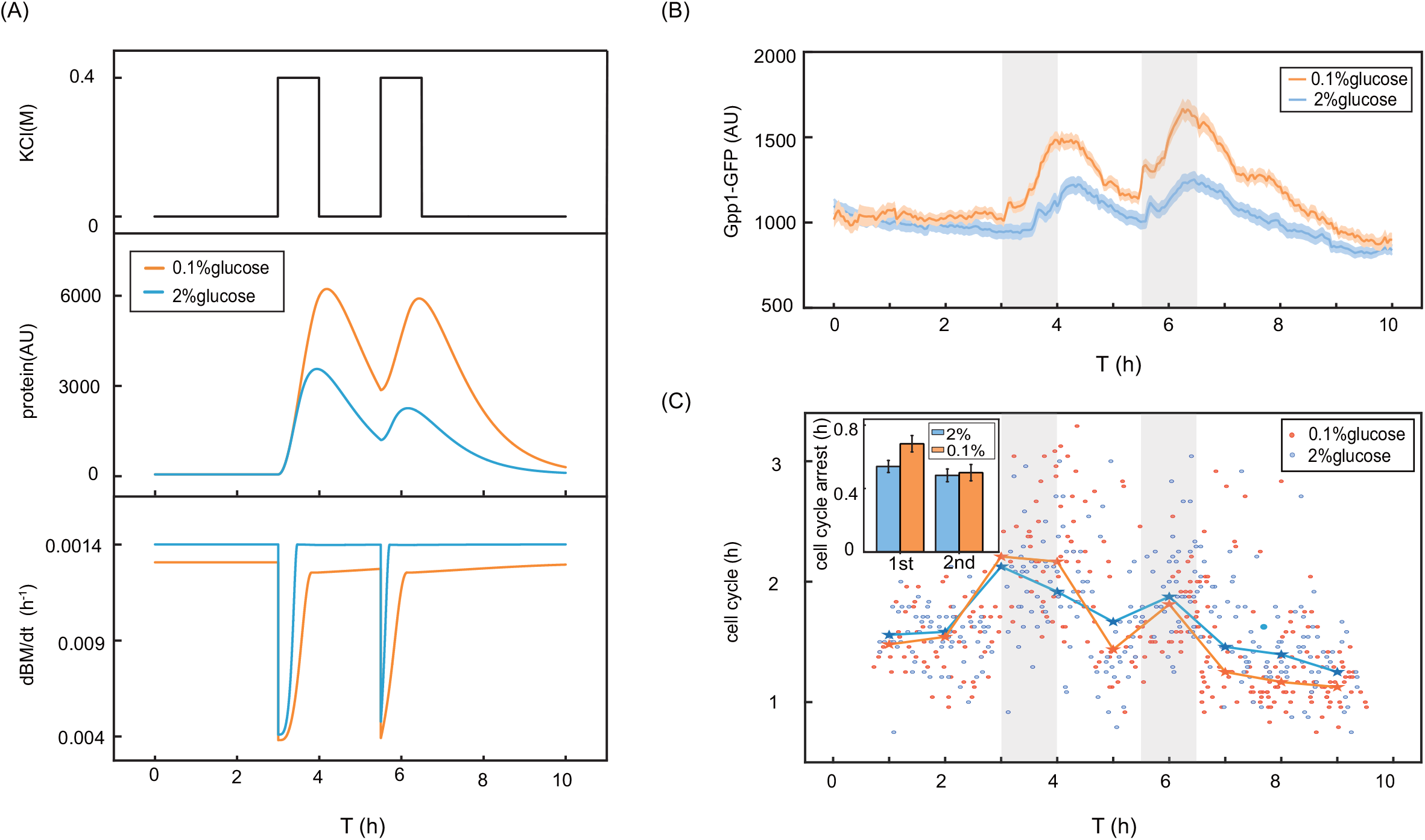
The enhanced response prepares for following potential stimuli. A. Model prediction of twice hyperosmotic stimulis. The stimulation lasted 1 hour, and then 1.5 h periods of normal medium was added, which is about a cell cycle long. Second hyperosmotic stimulation had the same period. B. Average Gpp1 expression to 0.4M periodic KCl stimulation. For cells in the glucose-limitation environment, due to the previous reponse cost, protein accumulation under the second stimulation was much higher than the first stimulation. C. Scattered points represent the cell cycle of single cells, and the x-axis is the middle moment of the two G1 phases. The dotted line indicates the median value of the single cell cycles in the previous half hour. Adaptation to the second osmostress input proceeded faster in both glucose concentrations. And due to accumulation of protein from the previous cycle, cells at 0.1% glucose accelerated recovery. Histogram inset is the statistics of cell cycle arrest time. There were 77 and 82 cells in 2, and 0.1% glucose concentrations respectively.

Collectively, these observations suggest that the enhanced response to hyper-osmotic stress, especially under low glucose conditions, can be helpful to adapt to the same following stress, which is manifested as a shortened recovery time. Excessive protein synthesis in a glucose limitation environment requires more tremendous cost. This strategy may be a selective advantage for cells since protein accumulation is a critical ability of microorganisms to adapt to potentially detrimental environmental fluctuations.

## Discussion

Environmental conditions shape the metabolism and gene regulation of all living cells. Currently, it is unclear how cells react to an adverse environmental condition when nutrient is limited. In this work, we systematically described protein dynamics of *S. cerevisiae* to hyperosmostress under different glucose environments. The reason for choosing this type of stress is because yeasts are often starved of carbon and suffered from dryness in nature. We observed differences in dynamic response profiles to varying levels of glucose concentration (Fig. 3A), and there was no significant difference in steady state growth rate (Fig. 2B). To understand what mechanisms involved in this phenomenon, we measured the nuclear translocation of general stress-responsive TF Msn2 and osmotic specific TF Hog1. Consistent with the synthesis rate inferred from downstream proteins, Hog1 significantly extended nuclear duration under low glucose conditions, while Msn2 increased slightly. Therefore, we found that under glucose limitation condition (0.1%), cells expressed much more osmotic-related proteins.

Considering the partition of limited proteome, it may be inferred from current research that there is no difference in resource allocation strategy when the growth rate is constant. However, we proved that even in a nutrient-limited environment where cell physiology was not significantly affected, cells had actually adopted a strategy to reduce ‘standby’ defense reserve. The stress response requires investment in the production of protective proteins and metabolites. Hence, cells have to compromise between the desire to grow and proliferate at a maximal rate and to prevent cellular damage under fluctuating conditions. Our results show that under glucose limitation environments, cells choose to optimize the biosynthesis and sacrifice reserve for rapid growth recovery from future perturbations (Fig. S9B). This ‘defense reserve saving’ strategy may prompt us to revise our view of how cells allocate resources.

Under abundant glucose conditions (>0.1%), carbon flux is redundant for growth and is stored in other metabolic branches. Upon osmoshock, this internal storage reroutes to glycerol production. Besides, cell cycle progression is arrested (Fig. S8A). According to our hypothesis, cells can adapt to osmostress and resume growth because of enough glycerol accumulation via protein synthesis and glycolytic reserves. Therefore, at a steady state, the inflow of glucose exceeds the efflux of glycerol synthesis, which enables cell cycle recovery. Nevertheless, at even lower glucose concentrations, more excess downstream proteins are nearly saturated for glycerol production, and the initial value of metabolic intermediate G3P is not enough to maintain sufficient glycerol. Therefore, cells still need to continuously route the glycolytic flux at steady state. Regarding the limited amount of glycolysis, in a poor nutrient environment, there is another trade-off between maximization of steady state exponential growth and perfect adaptation to osmostress. Cells can prioritize growth reduce defense costs, or use all resources to achieve stress response (Fig. S8B). Biologically, our work reveals a novel mechanism for yeast strategy under such condition. In a glucose limitation environment (<0.1%), biomass synthesis is restricted. After stimulated by hyperosmotic, Hog1 nuclear localization duration is prolonged and eventually can exit the nucleus, which means turgor pressure balance (Fig. S6). But the cell cycle cannot adapt to initial state (Fig. S9C). This implies that cells choose to sacrifice growth and preferentially meet the stress defense demands. Moreover, the more severe the stimulus, the greater the sacrifice for growth.

The second contribution of this work is to formulate an instance in which cells enhance stress response for faster recovery to future environmental changes (Fig. 6C). The reserve from sufficient nutrition can be on standby for faster adaptation to sudden changes. While under glucose limitation condition, more protein accumulation is required for adaptation, thus making the cell have a reserve for defense and a better response capacity to the same type of stimulation in the future. Therefore, in an environment where repeated or oscillating stress might be more probable, this strategy of enhanced response will be a fitness advantage.

In summary, our combined computational and experimental analysis reveal a mechanism that accounts for enhanced osmostress response under glucose limitation environments. We conclude that glycerol metabolism plays multiple roles in yeast adaptation to altered growth conditions, explaining the complex regulation of glycerol biosynthesis genes. Metabolic trade-off in poor nutrient adapts to the defense and biosynthetic demands of the cells. It can be inferred that yeast saves defense reserves, and once stimulated, it will require higher protein response costs and longer recovery time.

Our work focused on glucose limitation and hyper-osmotic stress. It is proved that at least in the case of osmostress under glucose limitation, yeast cells do not maximally mobilize their internal carbon sources to preserve for stress, thus sacrificing substantial speed in their recovery. How cells encode other stresses in limited nutrients remains unaddressed. For example, the response to oxidative or ethanol stress is closely connected with the activity of cAMP/PKA pathway and is involved in metabolism(Gutin *et al*, 2015b). When glucose is available, activation of the cAMP/PKA pathway stimulates glycolytic flux and suppresses stress response(Conrad *et al*, 2014). It would be interesting to investigate if it is a universal phenomenon that cells in a glucose limitation environment need an enhanced response for defense. In future studies, similar approaches might be used to discover similar metabolic flux distribution mechanisms. The apparent trade-off between growth and stress resistance is a potential source for the general growth strategies under different environmental conditions.

## Acknowledgements

This study was supported the National Key Research and Development Project (SQ2018YFA090070-03 and 2020YFA0906900) and the National Natural Science Foundation of China (11974002, 11774011, 12094054, and 11674010).

## Author Contributions

C.L. designed the study; W.S. and K.C. performed the experiments; C.L. and W.S. analyzed the data; W.S., Z.G., C.L. and Q.O. wrote the paper.

## Methods and materials

### Yeast strains

Yeast strains used in this study were from the S. cerevisiae GFP fusion library generated by Prof. Erin O’Shea and Prof. Jonathan Weissman (Huh *et al*, 2003). The yeast strains were generated from the BY4741 (MAT a his3Δ1 leu2Δ0 met15Δ0 ura3Δ0) strain background. To monitor cell cycle, we transferred MCM-mCherry plasmid #(pCT05) into yeast strains, which is provided by Tang Chao Laboratory(Yang *et al*, 2013).

Yeast cells were cultured overnight, then diluted into fresh medium and incubated for 4– 5 h at 30 °C to exponential growth (OD600_nm_ 0.4). Cells were then loaded into the microfluidic device and cultured at different glucose concentrations.

### Time-lapse microscopy experiments using a microfluidic platform

The microfluidic chip was mounted on the stage of a Nikon inverted microscope, with bright-field, GFP and m-Cherry images acquired every 2.5min using a 60× oil immersion objective. Full details of the microfluidic chip are given in the Supplementary Materials(Fig. S1)(Zhang *et al*, 2017).

Cells were maintained in synthetic drop out medium with amino acids and desired concentration of glucose (2%, 0.1% glucose). For glucose limitation medium, PBS buffer was added.

### Parameter characterization of stress response dynamics

In our characterization of the stress response, the abundance of protein is determined by the synthesis rate α and the decay rate β. Considering the timescale of fast response and slow protein degradation, the degradation rate can be ignored. Thus, decay term is considered to be the dilution rate of the protein. β is the volume change in a cell cycle, inversely proportional to the cell cycle T, and positively related to the volume ratio of daughter and mother cells when the cell budding V_m_/V_d_.

When cells are stimulated by osmostress, cell cycle arrest, decay rates decreased (Fig. S2B). After a cycle, decay rates were mostly restored. The cell cycle delay is related to the glucose concentration. So, it needs longer recovery time in low glucose environment.

We observed the single cell through the microfluidic platform to obtain the time sequence of the cell volume and the distribution during the budding of the cell, thereby obtaining the dilution term β. Thus, the simplified model captured key variables of cell response dynamics.

### Measuring Hog1 and Msn2 nuclear localization in single yeast cells

Localization of transcription factor Hog1 and Msn2 in a single cell was defined as the ratio of the average GFP pixel intensity in the estimated nucleus and the average GFP pixel intensity in the whole cell, as previously described (Chen *et al*, 2020b).

The amplitude of TF localization was quantified by measuring the single-cell TF localization signals that were above the threshold. The duration of Msn2 localization was quantified by measuring the average of all time intervals during which the single-cell Msn2 localization was above the threshold(Chatterjee & Acar, 2018).

